# MIMOSA: A model-independent framework for comparing transcription factor binding site motifs

**DOI:** 10.64898/2026.05.13.725009

**Authors:** Anton V. Tsukanov, Victor G. Levitsky

## Abstract

**Motivation:** Transcription factors (TFs) regulate gene expression by binding specific DNA sequences, which are commonly represented by motif models. Although position weight matrices (PWMs) remain the dominant motif representation, alternative models, such as Markov models, can capture interpositional dependencies and may provide higher predictive performance. Existing motif comparison tools, however, are designed mainly for PWMs or require reducing motifs to position weight matrices or position frequency matrices. This limits cross-model motif comparison, complicates interpretation of de novo motif discovery results, and hinders integration of diverse motif models into genomic analyses.

**Results:** We present MIMOSA (Model-Independent Motif Similarity Assessment), a model-independent framework for direct comparison of TF binding site (TFBS) motifs regardless of their mathematical representation. MIMOSA assesses motif similarity by comparing calibrated recognition profiles produced by different motif models on the same DNA sequence set, rather than by comparing motif parameters. In a cross-database benchmark on HOCOMOCO motifs, MIMOSA achieved retrieval performance comparable to established PWM-oriented tools, including Tomtom and MACRO-APE, with mean reciprocal rank (MRR) and Recall@*k* close to the best-performing methods. Pairwise ranking comparisons showed that MIMOSA captures a similarity signal consistent with existing approaches while providing a model-independent comparison strategy. Applying MIMOSA to de novo ATF3 ChIP-seq motifs showed that recognition-profile comparison distinguished alternative spacer variants represented as separate PWMs from their integration within more flexible models such as BaMM and Slim. Thus, MIMOSA enables quantitative cross-model motif comparison and supports interpretation of motif heterogeneity in TFBS analyses.

**Availability and implementation:** MIMOSA is implemented in Python and is freely available at https://github.com/ubercomrade/mimosa.

## Introduction

Transcription factors (TFs) regulate gene expression by binding specific sites in cis-regulatory DNA regions (Lambert et al., 2018). These transcription factor binding sites (TFBSs) are rarely identical in sequence; their variability is usually described by motifs, that is, statistical models of the sequence specificity of protein–DNA interactions (Wasserman and Sandelin, 2004). Motif structure reflects the properties of DNA-binding domains, TF dimerization, and the biophysics of protein–DNA contacts (Levitsky et al., 2025). The most widely used motif model is the position weight matrix (PWM), because it is straightforward to interpret. A classical PWM assumes positional independence: each position contributes additively to the site recognition score, which can be interpreted as an approximation of binding affinity (Berg and von Hippel, 1987).

However, many experimental and computational studies have shown that positions within binding sites can be interdependent (Benos et al., 2002; Bulyk et al., 2002; Slattery et al., 2014; Keilwagen and Grau, 2015; Kribelbauer et al., 2019; Cooper et al., 2023). Such dependencies can reflect non-additive protein–DNA contacts, DNA shape, TF dimerization, and cooperative binding of regulatory proteins (Amoutzias et al., 2008; Morgunova and Taipale, 2017). Thus, although the PWM remains a popular representation, more flexible alternative motif models can complement its predictions.

High-throughput methods such as ChIP-seq (Park, 2009), HT-SELEX (Jolma et al., 2010), and DAP-seq (O’Malley et al., 2016) have greatly expanded maps of TF binding specificity. These approaches generate DNA sequence sets enriched for TF binding sites, but the motifs themselves usually have to be reconstructed computationally. Researchers commonly reconstruct these motifs with de novo motif discovery methods, including STREME (Bailey, 2021) and HOMER (Heinz et al., 2010), which use the PWM model. PWM limitations have motivated methods that can account for more complex motif structure: Markov and Bayesian models, such as BaMM (Siebert and Söding, 2016) and InMoDe (Eggeling et al., 2017); discriminative approaches, such as DIMONT (Grau et al., 2013); dimeric motif models, such as MODER2 (Toivonen et al., 2020); and SiteGA, which is based on discriminant analysis and a genetic algorithm (Levitsky et al., 2007; Tsukanov et al., 2022). Neural network models, including DeepBind (Alipanahi et al., 2015) and later architectures (Chen et al., 2021; Wang et al., 2024), have also proved effective for TFBS prediction and for predicting sequence-level regulatory activity.

ChIP-seq analyses support the practical value of such models: SiteGA identified additional structural classes of binding sites of FoxA2 and SF1 TFs associated with distinct functional gene ontology terms (Levitsky et al., 2014, 2016). Another study combined the PWM, BaMM, and SiteGA motif models and increased the fraction of ChIP-seq peaks containing predicted sites for Arabidopsis thaliana TFs (Tsukanov et al., 2022). These results indicate that alternative motif models not only complement PWMs but can also reveal biologically meaningful classes of binding sites that are partially masked when a single matrix representation is used. Consequently, modern motif discovery is becoming multi-model: the same TF specificity can be represented by different mathematical objects, creating a need for their rigorous comparison.

After de novo motif discovery, a key step is to compare the discovered TFBS motifs with reference motif collections such as HOCOMOCO, JASPAR, and CIS-BP (Vorontsov et al., 2024; Ovek Baydar et al., 2026; Weirauch et al., 2014). For ChIP-seq data, this comparison determines whether a discovered motif corresponds to the target TF, a closely related TF from the same family, or a potential partner TF. This is particularly important for in vivo data, where motifs may reflect direct binding by the target TF or indirect binding through partner factors (Levitsky et al., 2019; Slattery et al., 2014). These databases also contain substantial TFBS motif redundancy from closely related TFs, differences between experimental systems, and differences in model construction (Ambrosini et al., 2020). This redundancy can also help describe TFBS motif variation within the same TF. Therefore, motif comparison, clustering, and annotation are required both for interpreting de novo motifs and for motif classification in large collections.

Existing motif comparison tools, including Tomtom (Gupta et al., 2007), STAMP (Mahony and Benos, 2007), MACRO-APE (Vorontsov et al., 2013), and MoSBAT (Santana-Garcia et al., 2022), are designed for PWMs and require converting a motif to a PWM. Such conversion may lose information about positional dependencies, dimeric structure, or other features of alternative motif models. Here, we ask whether two motif models recognize the same sequence specificity even when they have different internal representations. To address this issue, we introduce MIMOSA, a tool for comparing TFBS motifs by their functional output rather than by model parameters. MIMOSA treats motif models as functions that score DNA sequences and defines motif similarity by the concordance of these scores on a common sequence set. This framework places different motif architectures in a shared functional space, so comparisons depend on model behavior rather than mathematical representation or internal parameterization.

In this work, we benchmarked MIMOSA on the reference collection of PWM motifs and applied it to de novo motifs represented by diverse motif models, including PWMs, BaMMs, Slim, and DIMONT, recovered from an ATF3 ChIP-seq dataset Our results show that model-independent TFBS motif comparison facilitates motif annotation, identifies similar specificities across model classes, and extends the interpretation of structural diversity of TFBS motifs.

## Results

### A. Overview of the MIMOSA workflow

For each motif pair, MIMOSA first creates recognition profiles for a given set of DNA sequences. It then normalizes these profiles, enumerates pairwise shifts and orientations of the recognized motif signal, including forward and reverse-complement alignments, and selects the alignment that maximizes the chosen similarity metric (Figure 1). In this work, we used two metrics: cosine similarity, which characterizes similarity in the shape of local recognition profiles, and Dice similarity, which characterizes symmetric overlap of recognition signals (see Methods section). For database-scale annotation, MIMOSA also evaluates the statistical significance of the observed similarity against a precomputed pooled null distribution constructed from cross-class comparisons of shuffled HOCOMOCO motifs. The upper tail of this null distribution is approximated by a generalized extreme value distribution. Thus, MIMOSA returns a functional alignment of motif models together with a calibrated estimate of the statistical significance of their similarity, enabling comparison of motifs with different internal representations.

**Figure 1.**
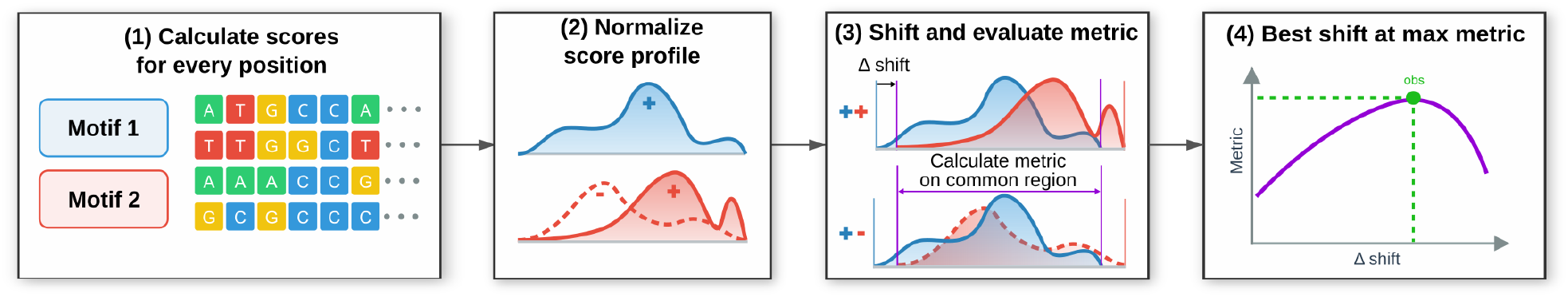
Schematic overview of the MIMOSA framework. Motif comparison based on recognition profiles. For each motif model, recognition scores are computed at all positions of a common sequence set, yielding positional recognition profiles. Raw scores are transformed into calibrated values, for example −log_10_(FPR), where the false-positive rate (FPR) is estimated from the corresponding background score distribution. Pairwise similarity between motif models is computed after shifting one profile relative to the other over all allowed offsets and orientation combinations. The similarity metric is evaluated on the common aligned region for each shift, and the maximum value over all shifts and orientations defines the optimal functional alignment of the motif models.

### B. Cross-database benchmark against established motif comparison methods

To compare MIMOSA with established tools, we used HOCOMOCO motifs: in vitro-derived motifs as queries and in vivo-derived motifs as targets. This cross-experimental design evaluates the ability of a method to identify motifs corresponding to the same TF independently of the experimental source. We evaluated MIMOSA in two variants, mimosa-dice and mimosa-cosine, and compared its results with those from Tomtom, STAMP, MACRO-APE, and MoSBAT.

We first assessed whether different tools produced concordant rankings of target motifs. For each pair of tools, we calculated Kendall’s rank correlation coefficient *τ* between the estimates of the corresponding similarity metrics. For MIMOSA, we used Dice similarity and cosine similarity; for MACRO-APE, the Jaccard index; for Tomtom, −log_10_(*E*-value); and for STAMP and MoSBAT, −log_10_(*P*-value). All pairwise correlations were positive, but most values were only moderately high (*τ ≈* 0.3–0.5; Figure 2). This indicates a shared ranking signal across tools. At the same time, correlations among established non-MIMOSA tools were also moderate, showing that MIMOSA is not an outlier and that the full ordering of target motifs, particularly weak matches to a query motif, depends substantially on the similarity function used.

**Figure 2.**
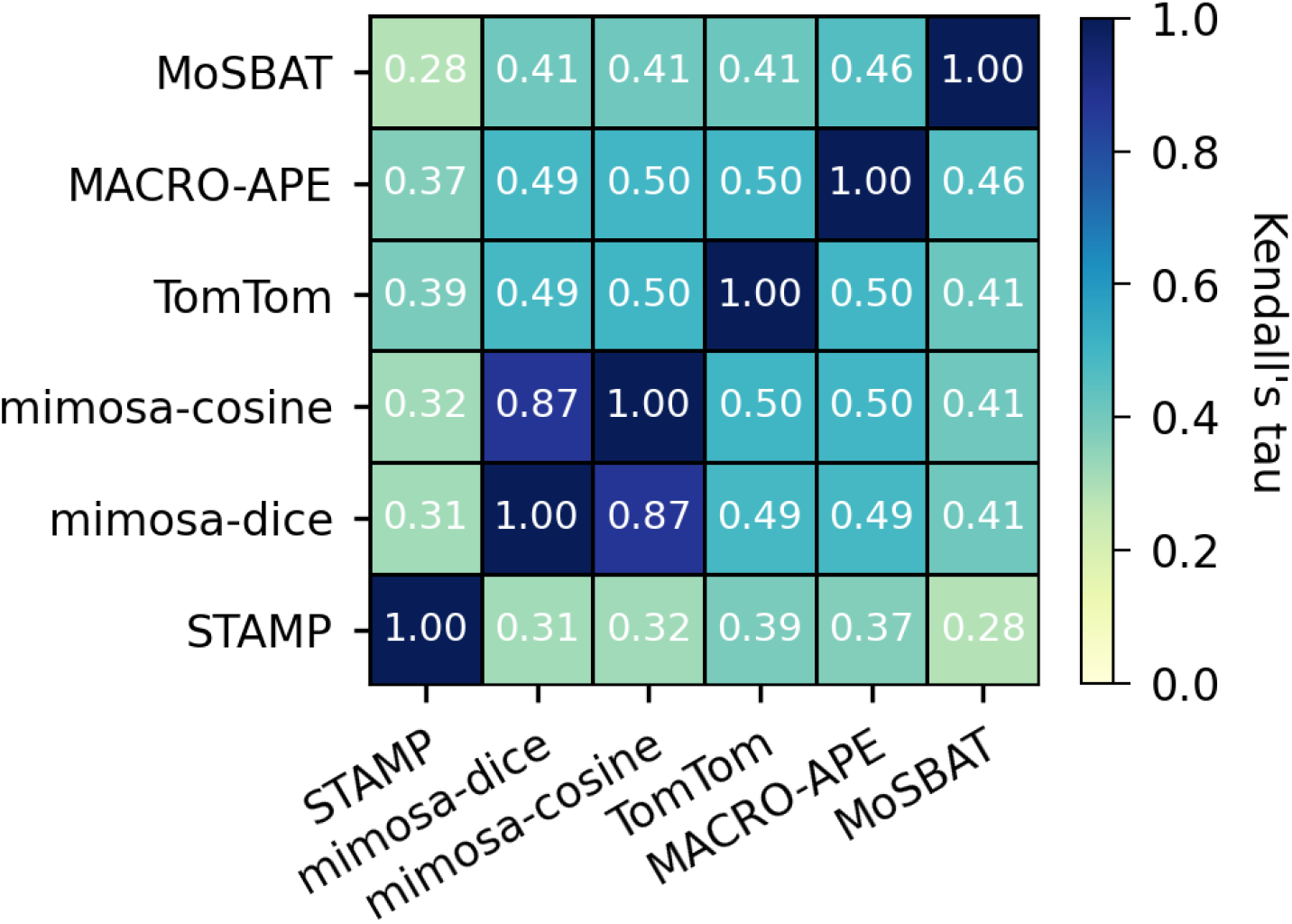
Pairwise concordance of motif rankings across comparison tools. The heatmap shows pairwise Kendall rank correlations (*τ*) between mimosa-dice, mimosa-cosine, MACRO-APE, Tomtom, STAMP, and MoSBAT. Higher values indicate stronger similarity between rankings of candidate target motifs in the benchmark set.

Thus, pairwise correlations indicate only limited concordance of complete motif rankings among tools. However, this test does not show how highly each tool ranks the truly relevant motif among all targets. We therefore evaluated retrieval accuracy using the mean reciprocal rank (MRR) and Recall@*k* at *k* = 1, 5, and 10 (Figure 3). Recall@*k* denotes the fraction of queries for which a correct motif is found among the top *k* predictions. A target motif was considered correct if it was annotated to the same TF as the corresponding query motif.

**Figure 3.**
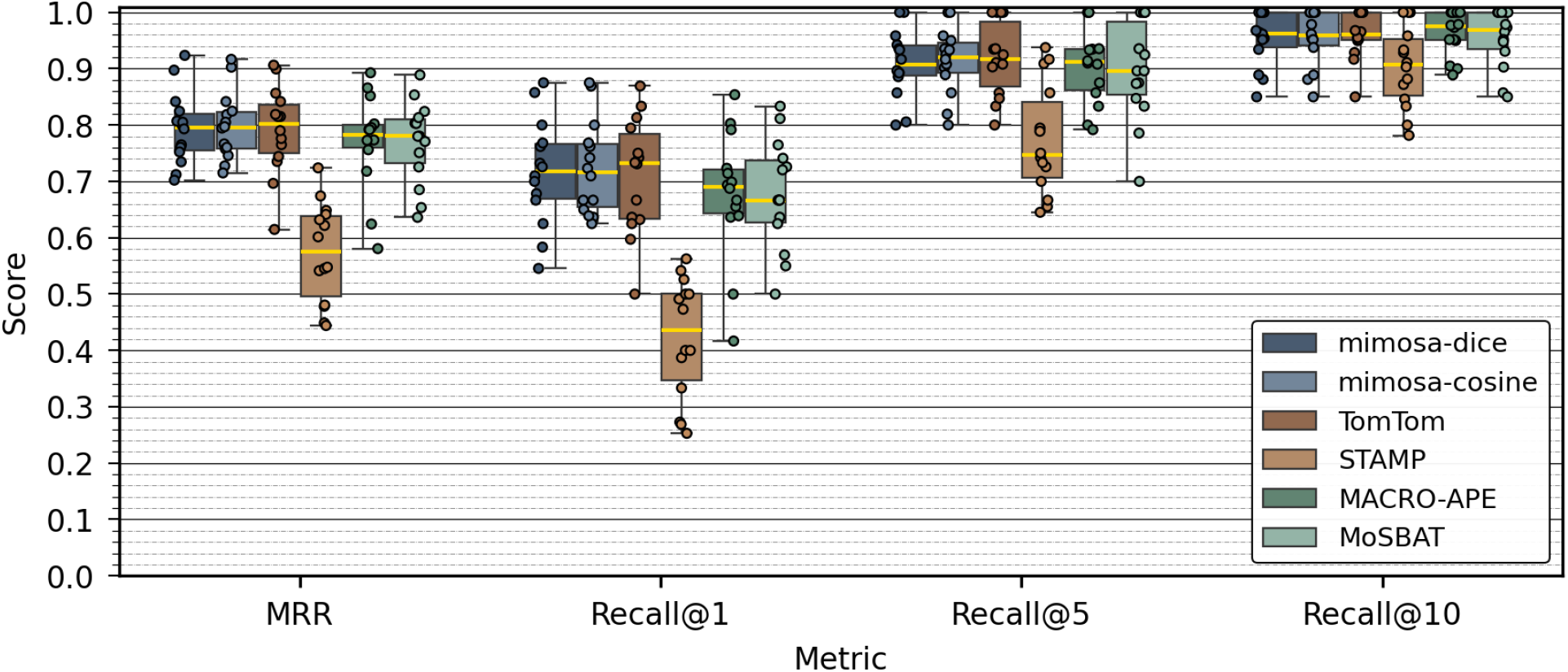
Retrieval performance of MIMOSA and established motif comparison tools. Ranking quality was evaluated using mean reciprocal rank (MRR) and Recall@*k* at *k* = 1, 5, and 10. The x-axis shows the ranking metrics, and individual points show values for separate transcription factor classes. A target motif was considered a correct match if it was annotated to the same TF as the corresponding query motif. The comparison includes mimosa-dice, mimosa-cosine, Tomtom, STAMP, MACRO-APE, and MoSBAT.

Tomtom obtained the highest mean MRR (0.833), followed by mimosa-cosine (0.826), mimosa-dice (0.825), and MACRO-APE (0.823). Differences among these tools were small, whereas MoSBAT showed a slightly lower but still comparable MRR (0.808). Recall@*k* showed a similar pattern: Tomtom achieved Recall@1 = 0.773 and Recall@5 = 0.915, whereas MIMOSA performed similarly. For MIMOSA, Recall@1 was 0.765 for cosine and 0.766 for Dice; Recall@5 was 0.903 for cosine and 0.911 for Dice. MACRO-APE also showed comparable accuracy and reached the highest Recall@10 among the methods tested (0.954). In contrast, STAMP performed substantially worse than the other tools, with MRR = 0.586 and Recall@1 = 0.444, although its recall increased markedly when the search window was expanded to the top 5 and top 10 positions (Recall@5 = 0.762 and Recall@10 = 0.876).

A class-wise analysis showed that retrieval performance strongly depended on the DNA-binding domain class of the TF (Supplementary Figures S3–S6). All methods most consistently identified motifs of Rel homology region (RHR) factors and C2H2 zinc finger factors, with approximate MRR values of 0.90–0.92 and Recall@1 values of 0.83–0.88. MIMOSA performed particularly well for high-mobility group (HMG) domain factors, other C4 zinc finger-type factors, paired box factors, and RHR factors. Tomtom retained an advantage for homeodomain factors, basic helix-loop-helix (bHLH) factors, and T-box factors. For several classes, including SMAD/NF-1 DNA-binding domain factors, Grainyhead domain factors, bZIP factors, and T-box factors, exact top-rank identification was less stable: Recall@1 decreased, whereas Recall@5 and Recall@10 often remained high. This indicates that in more complex or less clearly separable TF classes, the correct motif often remains among the closest candidates but does not always occupy the first rank.

Together, these results show that both MIMOSA variants achieve retrieval accuracy comparable to the strongest established motif comparison tools. Tomtom retained a small advantage in mean MRR and Recall@1, but the gap between Tomtom and MIMOSA was narrow. This was particularly evident when the top several ranks were considered: Recall@5 and Recall@10 values for MIMOSA, Tomtom, MACRO-APE, and MoSBAT became similar. Thus, comparison based on recognition profiles can effectively recover biologically relevant correspondences between motifs while preserving the key advantage of MIMOSA: the ability to compare motifs represented by different model classes.

### C. Cross-model analysis of ATF3 TFBS motifs from ChIP-seq data

To test how MIMOSA can support interpretation of de novo motif discovery results obtained with different motif-modeling approaches, we analyzed ATF3 TFBS motifs derived from a ChIP-seq dataset (GTRD ID PEAKS037311). All de novo motif discovery tools used in this analysis identified ATF3-like motifs with a characteristic AP-1/CRE-like dimeric organization comprising two half-sites, TGA and TCA (Figure 4A). STREME returned two distinct PWM motifs, PWM-1 and PWM-2, with the consensus sequences TGAnTCA and TGAnnTCA, respectively, which differ by one nucleotide in spacer length between the half-sites (Figure 4A). In contrast, dependency-aware models, especially BaMM and Slim, represented both spacer variants within a single motif. The DepLogo representations show this pattern (Figure 4A), with positional dependencies and signal heterogeneity within the motif; the site-alignment blocks in the Supplementary Materials also show that a single motif contains groups of sequences corresponding to the 1-bp and 2-bp spacer variants (Supplementary Figure S7). DIMONT recovered a core motif with partial ATF3-like similarity, but its representation was more distinct from those of the other tools.

**Figure 4.**
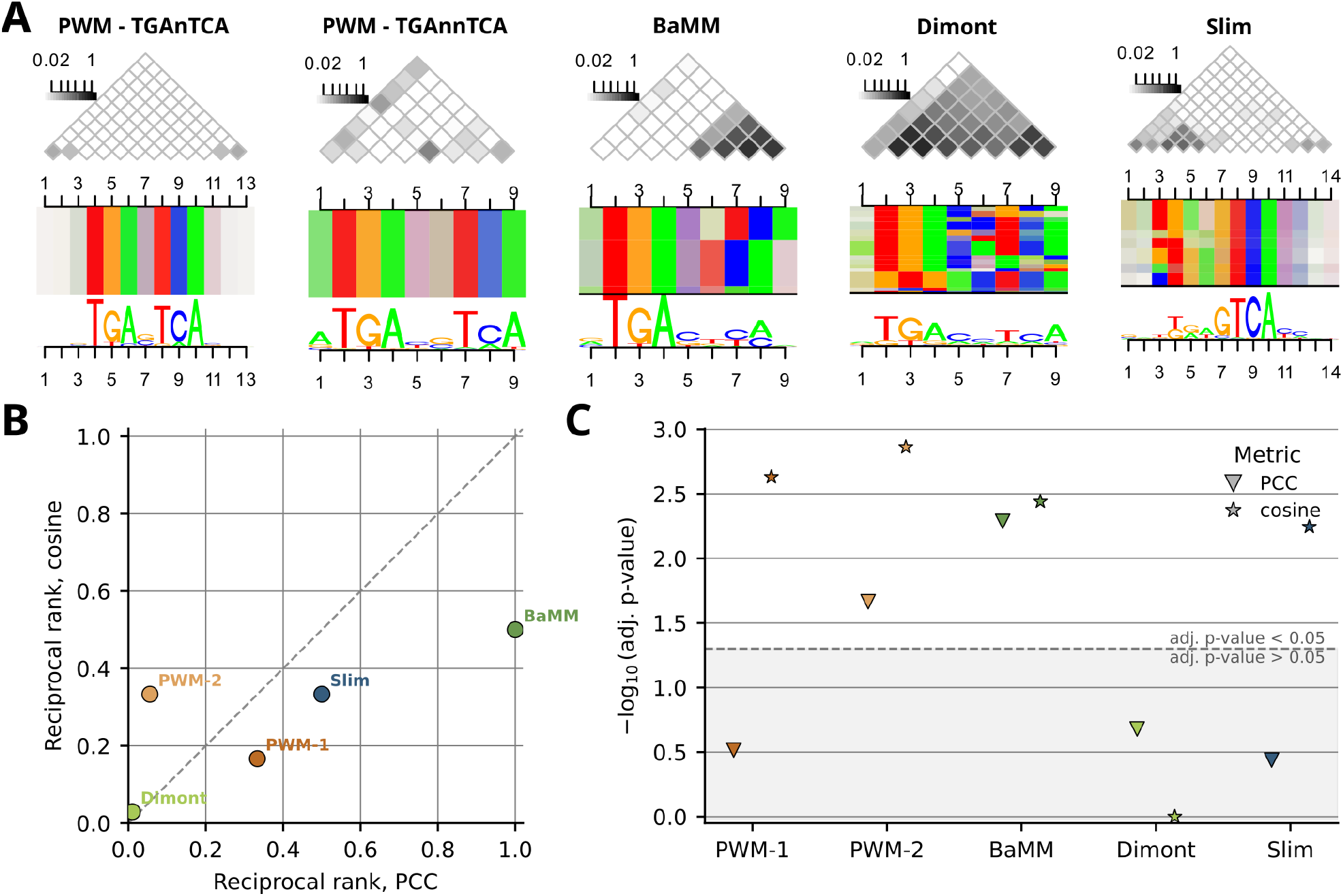
DepLogo representation and HOCOMOCO annotation of ATF3 motifs recovered de novo from ChIP-seq data. (A) DepLogo representations for five motifs obtained from the ATF3 ChIP-seq dataset: two PWM motifs with the consensus sequences TGAnTCA and TGAnnTCA, and motifs produced by BaMM, DIMONT, and Slim. For each motif, the lower panel shows the classical sequence logo, the central panel shows the site block representation, and the upper triangular matrix shows pairwise positional dependencies; grayscale intensity indicates dependency strength. (B) Reciprocal rank of the best ATF3 HOCOMOCO match for each recovered motif under profile-based cosine similarity and matrix-based Pearson correlation coefficient (PCC) between reconstructed position frequency matrices (PFMs). Higher reciprocal rank indicates a stronger ATF3 database match; the diagonal line marks equal ranks under the two strategies. (C) Statistical significance of the best ATF3 HOCOMOCO match for each recovered motif, shown as −log_10_ adjusted P-value. Stars denote profile-based cosine similarity, triangles denote matrix-based PCC, and the dashed line marks adjusted P = 0.05.

To confirm that the recovered de novo motifs corresponded to the expected ATF3 specificity, we compared them with HOCOMOCO database motifs using two complementary strategies (Figure 4B,C). We treated profile-based cosine similarity as the primary annotation metric because it compares motif recognition profiles directly and therefore preserves the behavior of motifs represented by different model classes. In contrast, the Pearson correlation coefficient (PCC) was computed after reconstructing position frequency matrices (PFMs) from high-scoring TFBSs predicted in ChIP-seq peaks at the threshold −log_10_(FPR) *≥* 3.0. PCC therefore represents a matrix-based projection of the original models. For heterogeneous or dependency-aware models, such a projection can average alternative site variants and reduce positional information at individual positions, as also visible in the sequence logos.

Profile-based database matching placed PWM-1, PWM-2, BaMM, and Slim within the bZIP/AP-1/CRE-like motif space. The best HOCOMOCO matches were NFE2 for PWM-1, ATF2 for PWM-2, JUN for BaMM, and JUN for Slim, and ATF3 motifs appeared among the highest-ranked matches for these four models (ranks 6, 3, 2, and 3, respectively). These ATF3 matches were significant after FDR correction (Figure 4C). DIMONT showed a distinct annotation pattern: its top profile-based match was the nuclear receptor motif RXRG, and the best ATF3 match was ranked only 34th and was not statistically significant. PCC-based annotation ranked ATF3 first for BaMM, second for Slim, third for PWM-1, 18th for PWM-2, and 98th for DIMONT. Among these ATF3 matches, only BaMM and PWM-2 passed FDR correction. Thus, HOCOMOCO annotation supported the expected ATF3/AP-1-like specificity of the PWM, BaMM, and Slim models, whereas DIMONT appeared to capture a more divergent signal.

We used two approaches to compare these five motifs pairwise (Figure 5A). First, we compared recognition profiles using cosine similarity. The profile-based graph connected BaMM to both ATF3 PWM motifs, with similarity values of 0.88 for PWM-1/BaMM and 0.86 for PWM-2/BaMM. Slim showed the same pattern, with similarity values of 0.82 for PWM-1/Slim and 0.77 for PWM-2/Slim, and BaMM and Slim were also closely connected (0.89). These results support the hypothesis that BaMM and Slim recognize the 1-bp and 2-bp spacer variants of the ATF3 site within a single motif. The absence of a direct edge between PWM-1 and PWM-2 is consistent with their interpretation as separate spacer variants. DIMONT did not connect to the main AP-1/CRE-like cluster in the profile-based graph, indicating more distinct recognition behavior.

**Figure 5.**
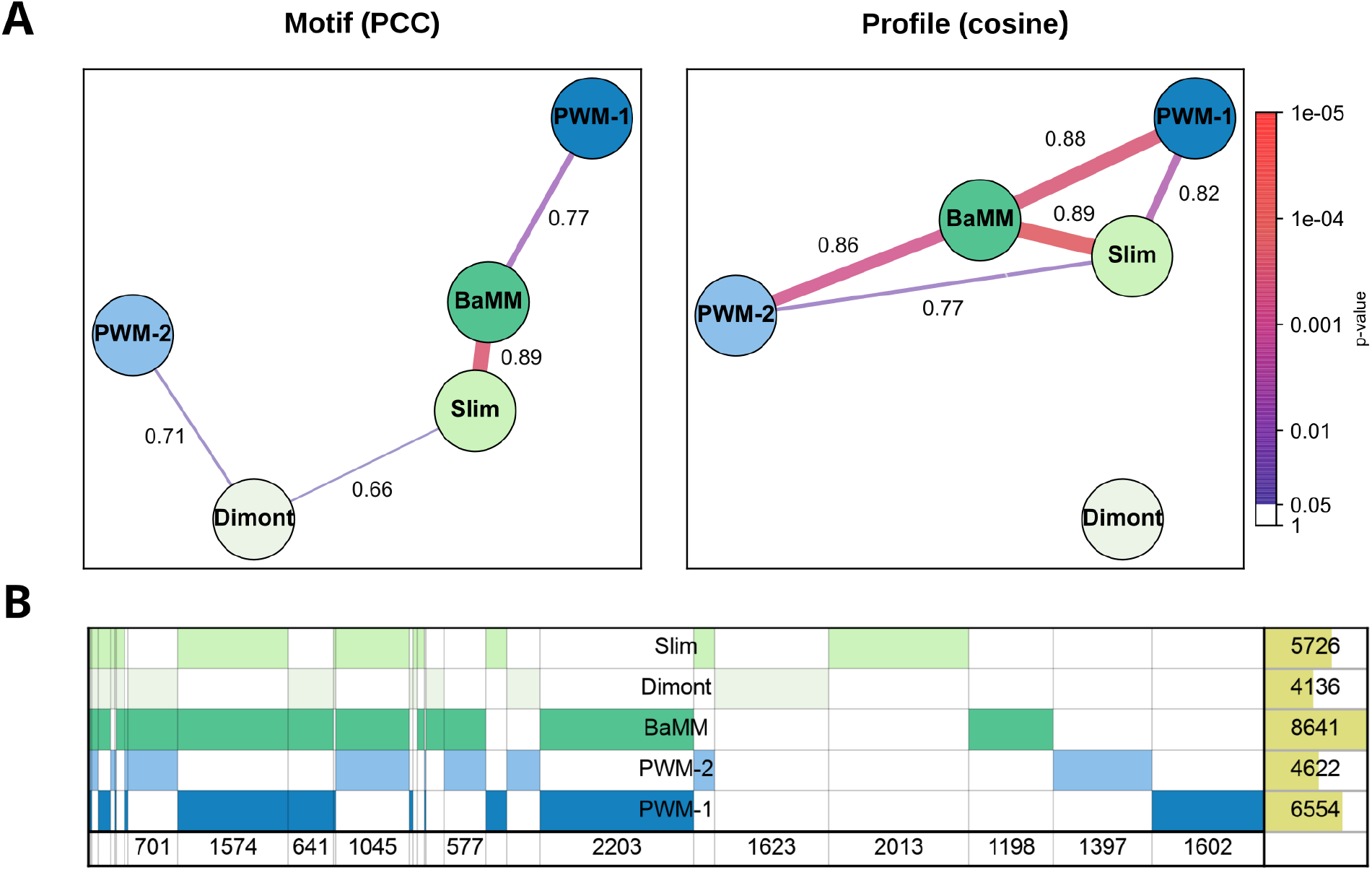
(A) Pairwise similarity of ATF3 motifs recovered de novo from ChIP-seq data. Network representation of pairwise motif similarity estimated by two approaches: matrix-based comparison of reconstructed position frequency matrices (PFMs) using Pearson correlation coefficient (PCC; Motif, PCC) and recognition-profile comparison using cosine similarity (Profile, cosine). Edge labels show similarity values, and edge color indicates the corresponding P-value. Higher similarity values indicate greater motif similarity. Profile-based cosine compares model recognition behavior directly, whereas matrix-based PCC summarizes each model through a single reconstructed PFM and can therefore reduce positional information for heterogeneous or dependency-aware motifs. (B) SuperVenn diagram of overlaps between predicted TFBS sets for ATF3 motif models. Horizontal bars correspond to site sets predicted by BaMM, Slim, DIMONT, and the two PWM motifs. Values on the right show the total number of sites predicted by each model, and values at the bottom show the sizes of the corresponding intersections between sets. At the selected score threshold, PWM-1 and PWM-2 have no overlapping predicted sites, indicating separation of the two spacer variants of the ATF3-like motif. Overlaps between BaMM and both PWM motifs indicate that BaMM recognizes sites corresponding to both spacer variants. The recognition threshold was −log_10_(FPR) *≥* 3.0, where FPR denotes the false-positive rate.

Second, we performed a matrix-based motif comparison. For each model, we selected candidate binding sites from high-scoring predictions, reconstructed PFMs from these sites, and compared the resulting matrices using PCC. The PCC graph showed a different topology: BaMM was connected to PWM-1 (0.77) and Slim (0.89), whereas DIMONT was connected to PWM-2 (0.71) and Slim (0.66). Thus, profile-based and matrix-based comparisons reflected different patterns of similarity. The former assesses functional overlap between motif predictions irrespective of the motif model, whereas the latter summarizes each model through a single reconstructed PFM and can skew similarity relationships for non-PWM motifs.

Overall, these results show that direct profile-based comparison is better suited for cross-model motif annotation than matrix-based comparison after PFM reconstruction. For dependency-aware models, PFM projection can average alternative site variants and reduce position-specific information, whereas profile-based similarity preserves the recognition behavior of the original models. Thus, MIMOSA confirmed the expected ATF3/AP-1-like specificity for PWM, BaMM, and Slim motifs and identified DIMONT as a more divergent case. However, pairwise similarity graphs alone do not show how the corresponding site predictions are partitioned among models. We therefore complemented the similarity analysis with an overlap analysis of predicted binding sites.

Site-level overlap and anchor-based profile alignment. To determine whether the model-level similarity patterns were reflected at the level of individual predictions, we next analyzed overlaps between predicted binding sites. At the selected score threshold, the SuperVenn analysis showed that the two PWM models identified non-overlapping site sets, whereas BaMM overlapped with both spacer variants (Figure 5B). The absence of overlap between PWM-1 and PWM-2 is consistent with their lower profile-based and matrix-based similarity. At the same time, BaMM overlapped with both PWM motifs and predicted the largest number of sites among all models: 8,641 sites for BaMM, 6,554 for PWM-1, 5,726 for Slim, 4,622 for PWM-2, and 4,136 for DIMONT. Together, these results show that MIMOSA can quantify similarity between motif models of different architectures and distinguish two situations: separation of alternative spacer variants into distinct PWM motifs and integration of these variants into a more flexible motif model.

To explain these site-level and model-level differences directly, we compared mean recognition profiles around top-scoring predicted sites using anchor-based alignment (Figure 6A–C).

**Figure 6.**
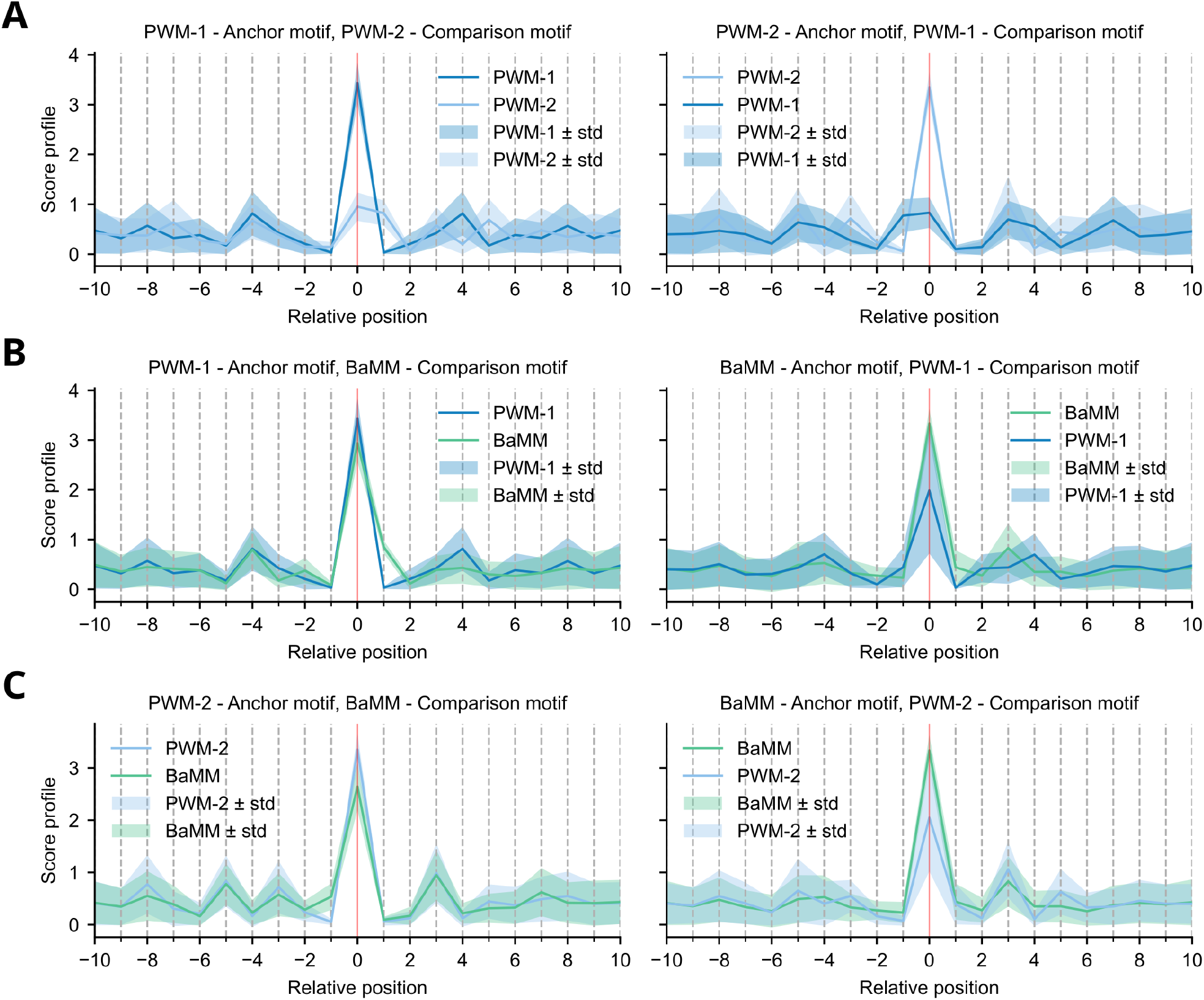
Anchor-based alignment of ATF3 motif recognition profiles. Pairwise alignments of recognition profiles are shown for (A) PWM-1/PWM-2, (B) PWM-1/BaMM, and (C) PWM-2/BaMM. In each row, the two panels show reciprocal anchoring schemes: the left panel is centered on high-scoring sites of the first motif model, whereas the right panel is centered on high-scoring sites of the second motif model. Profiles were aligned using the optimal MIMOSA shift obtained with cosine similarity on foreground sequences. The vertical line indicates the anchor position at relative coordinate 0; dashed lines mark flanking positions within the *±*10 bp window. Solid lines show the mean score profile across selected windows, and shaded areas show the mean *±* standard deviation. Strong agreement between aligned profiles indicates recognition of similar positions, whereas differences in peak height and variability reflect model-specific recognition behavior. Anchor sites were selected using the threshold −log_10_(FPR) *≥* 3.0, where FPR denotes the false-positive rate.

The anchor-based alignments directly explained the observed similarity and overlap patterns. For each motif pair, we examined both reciprocal anchoring directions. For the PWM-1/PWM-2 pair, sites anchored by one PWM motif had low scores for the other motif, and vice versa, indicating that the two PWMs capture different spacer variants. The pattern differed for BaMM: around anchor sites from both PWM-1 and PWM-2, BaMM showed a high mean signal with relatively low variability, indicating that it robustly recognizes both ATF3 spacer variants. However, when BaMM was used as the anchor model, the PWM-1 and PWM-2 profiles showed increased variability: each PWM model scored only the subset of BaMM predictions corresponding to its own spacer variant well, whereas the remaining subset was weak for that PWM model. This explains why BaMM has high profile similarity to both PWM models but can be closer to only one reconstructed PWM representation in the matrix-based comparison.

## Discussion

Comparison of TFBS motifs remains a key step in interpreting de novo motif discovery results from ChIP-seq and other high-throughput experiments that characterize TF–DNA interactions. In practice, researchers typically obtain a set of enriched motifs and must compare them with known binding site models from databases such as HOCOMOCO, JASPAR, and CIS-BP (Vorontsov et al., 2024; Ovek Baydar et al., 2026; Weirauch et al., 2014). This comparison is required both for annotating motifs that correspond to the target TF and for identifying motifs of potential partner factors that participate in composite regulatory elements.

Most motif comparison methods have focused on traditional PWMs. The PWM model remains popular because it is interpretable, convenient, and readily visualized with sequence logos (Schneider and Stephens, 1990). Many specialized tools have therefore been developed for comparing PWM motifs (Gupta et al., 2007; Mahony and Benos, 2007).

However, natural TFBSs can show positional dependencies, variable flanking contexts, differences in site affinity, and alternative structural variants of the same motif. Modern ChIP-seq, HT-SELEX, and related sequencing-based analyses show that binding sites can be more complex than traditional PWM models assume. One reason is that transcriptional regulation often involves cooperative action of multiple TFs (Slattery et al., 2014; Keilwagen and Grau, 2015; Levo et al., 2015; Elmas et al., 2017; Omidi et al., 2017; Kribelbauer et al., 2019; Castellanos et al., 2020; Tsukanov et al., 2022). This helps explain TFBS motif variation within the same TF in large motif collections.

This complexity is particularly important for motifs derived from in vivo data. TFs often function within protein complexes, and interactions between TFs can produce composite or overlapping motifs (Amoutzias et al., 2008; Morgunova and Taipale, 2017; Levitsky et al., 2019; Muñoz et al., 2025). In addition, the same TF can bind with different partners in different cellular contexts (Pihlajamaa et al., 2014), and a substantial fraction of functional sites may have low or intermediate affinity (Kribelbauer et al., 2019; Muñoz et al., 2025). Previous studies have shown that more flexible models, such as BaMM, SiteGA, and other alternative motif models, can identify additional classes of binding sites that are not fully captured by PWMs (Levitsky et al., 2014, 2016; Tsukanov et al., 2021, 2022). Such models can also be visualized: for example, dependency logos can display positional dependencies and site heterogeneity without reducing a motif to a simple PWM structure (Grau et al., 2019).

However, the development of alternative motif models creates a methodological problem: motifs represented by different mathematical formalisms are difficult to compare directly. The internal parameters of PWMs, BaMMs, Slim, DIMONT, or other models have different meanings and cannot always be converted to a common matrix representation without information loss. Thus, comparison based solely on internal model parameters limits the interpretation of de novo motif discovery results, especially when different tools describe the same biological signal in different ways.

Here, we present MIMOSA, a framework that addresses this problem by shifting motif comparison from model parameters to functional behavior. MIMOSA does not directly compare the internal parameters of motif models. Instead, it projects each model into a common functional space: the model scores the same set of DNA sequences, and MIMOSA calibrates and compares the resulting recognition profiles. This strategy makes it possible to compare motifs represented by different model classes, provided that each model can assign scores to sequences.

The benchmark analysis on HOCOMOCO motifs showed that MIMOSA achieves retrieval performance comparable to the strongest established motif comparison tools. In the cross-experimental setting, where in vitro motifs were used as queries and in vivo motifs were used as targets, both MIMOSA variants showed MRR and Recall@*k* values close to those of Tomtom and MACRO-APE. Tomtom retained a small advantage in mean MRR and Recall@1, whereas MACRO-APE achieved the highest recall when the search window was expanded to ten ranks. Nevertheless, differences between Tomtom, MACRO-APE, and the two MIMOSA variants were small, whereas STAMP was clearly less accurate in recovering the first correct match.

This result is important not because MIMOSA universally outperforms specialized PWM tools, but because it maintains competitive retrieval accuracy in a more general setting. Tomtom and MACRO-APE are optimized for comparing motifs in matrix representation, whereas MIMOSA uses only model recognition profiles. Therefore, the high accuracy of MIMOSA shows that functional comparison of recognition profiles preserves biologically relevant similarity signals between motifs while avoiding restriction to PWM representations.

Pairwise correlations among tool rankings further show that motif comparison does not reduce to a single universal ordering of target motifs. All methods showed positive correlations, but Kendall’s *τ* remained moderate for most tool pairs. This means that different similarity functions emphasize different aspects of a motif, especially among weak or ambiguous candidates. Thus, global concordance of rankings is not by itself a sufficient measure of tool quality. From a practical perspective, the more important question is how highly a tool places the biologically correct motif near the top of the list, as captured by MRR and Recall@*k*.

The class-wise analysis also showed that task difficulty depends on the DNA-binding domain class of the TF. For some classes, such as Rel homology region factors and C2H2 zinc finger factors, correct correspondences were recovered robustly by almost all methods. For other classes, including SMAD/NF-1 DNA-binding domain factors, Grainyhead domain factors, bZIP factors, and T-box factors, the first rank was less stable, although the correct motif often remained among the closest candidates. This indicates that within some TF families, boundaries between related motifs can be diffuse, and biologically relevant correspondences are not always unambiguously defined by a single top-ranked hit.

The ATF3 cross-model analysis provides the most illustrative example of the advantages of MIMOSA. All de novo tools identified an ATF3-like AP-1/CRE motif with a dimeric organization, but different models represented this signal differently. STREME split it into two PWM motifs differing in spacer length between the half-sites. In contrast, BaMM and Slim represented both spacer variants within a single motif, as observed both in DepLogo representations and in the distribution of sites.

Recognition-profile comparison showed that BaMM and Slim are functionally close to both ATF3 PWM variants. At the same time, the two PWM motifs were substantially less similar to each other, consistent with their interpretation as two alternative spacer variants. In other words, profile-based comparison revealed that more flexible models can recognize both spacer variants of the ATF3 site, whereas each separate PWM describes only one of them. This conclusion was further supported by the analysis of anchor-based recognition profiles and overlaps between predicted binding sites: BaMM overlapped with both PWM motifs, whereas PWM-1 and PWM-2 showed no overlap at the selected threshold.

Importantly, profile-based and matrix-based comparisons in this example produced complementary rather than identical results. Comparing reconstructed PFMs using PCC was sensitive to which spacer variant of high-scoring sites dominated matrix construction. Thus, BaMM, Slim, or DIMONT could appear closer to one PWM variant even when their recognition behavior overlapped functionally with both. This distinction highlights a central property of MIMOSA: recognition profiles reflect not only the shape of an averaged motif but also the behavior of the model across many sequences. Therefore, MIMOSA can distinguish two situations that may look similar in visual logo analysis: separation of alternative spacer variants into distinct PWM motifs and integration of these variants into a more flexible motif model.

Together, the benchmark analysis and the ATF3 case study show that MIMOSA is useful not only as a quantitative motif comparison tool but also as an aid for interpreting de novo motif discovery results. It enables comparison of motifs across architectures, identifies functional overlap between models, and distinguishes partial matrix similarity from similarity in predictive behavior. This is particularly important for ChIP-seq analysis, where the same biological signal can be represented by several motifs differing in length, spacer, positional dependencies, or the set of recognized sites.

Despite these advantages, the approach has limitations. First, profile similarity depends on the sequence set used to compute recognition profiles. If the selected sequence set poorly represents relevant site variants, model similarity can be underestimated or overestimated. Second, results depend on score calibration, the background distribution, and selected thresholds, especially when overlaps between predicted TFBSs are analyzed. Finally, the detailed ATF3 analysis represents one illustrative case; additional case studies across TF families and motif architectures are needed to assess how widespread such situations are.

MIMOSA can be extended in several directions. First, it will be useful to systematically examine how the choice of foreground and background sequences affects the robustness of profile similarity. Second, MIMOSA can be applied to a broader range of alternative motif models, including deep neural network models, provided that positional recognition profiles can be derived from them. Third, profile-based comparison can be used not only for annotation of de novo motifs but also for clustering motifs within large collections, identifying alternative structural variants, and analyzing evolutionary conservation of recognition patterns.

Overall, our results show that motif comparison through recognition profiles is an effective and interpretable alternative to direct comparison of model parameters. MIMOSA achieves accuracy comparable to strong specialized methods for PWM motif comparison while removing the restriction on the internal representation of a motif. This makes MIMOSA particularly useful for modern TFBS analysis, where classical PWMs, dependency-aware models, and other scoring frameworks need to be considered jointly in a common functional space.

## Methods

### D. Overview of the MIMOSA profile framework

Profile construction and normalization. Let *M*_1_ and *M*_2_ denote two motif models to be compared, and let *S* be a common set of sequences. For each model, MIMOSA scans the sequences in the forward and reverse-complement orientations and constructs two positional recognition profiles, 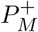 and 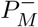. In each profile, rows correspond to individual sequences and columns correspond to possible motif start positions.

Because different model types can use incomparable score scales, profiles are normalized separately for each model. For this purpose, MIMOSA builds an empirical distribution of positional scores for each model using a calibration sequence set. For each raw score *r*, MIMOSA first estimates the empirical false-positive rate

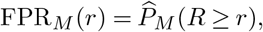

that is, the probability of obtaining a score not lower than *r* for the same model on the calibration set. The score is then transformed to a logarithmic scale:

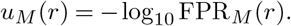

Thus, *u*_*M*_ (*r*) is an empirical log-FPR score for a position. It reflects how rare a raw score at this position is relative to the model-specific background score distribution rather than the absolute scale of the original model score. For example, *u*_*M*_ (*r*) = 2 corresponds to FPR = 10^−2^, whereas *u*_*M*_ (*r*) = 3 corresponds to FPR = 10^−3^.

Selection of informative windows. Next, MIMOSA compares local windows around the most informative signals on profiles. For a pairwise comparison, the same threshold *T* on the empirical log-FPR scale is applied to both profiles. A position is considered an anchor if its normalized score satisfies *u*_*M*_ (*r*) *≥ T*. This threshold selects regions in which the model recognizes a motif-like signal. Within the selected windows, the original continuous values *u*_*M*_ (*r*) are retained and then used to compute Dice and cosine similarity (see below). Therefore, a higher threshold makes the comparison more focused on stronger signals, whereas a lower threshold includes weaker signals, including potentially spurious predictions that may not correspond to true TF binding.

Orientations and shifts. For each motif pair, MIMOSA considers four combinations of motif profile orientations: ++, *−−*, +−, and −+. The first symbol refers to the profile of the first model and the second symbol to the profile of the second model. The ++ and *−−* combinations compare motifs on the same strand, using forward or reverse-complement profiles, whereas +−and −+ compare opposite-strand orientations.

For each orientation, integer shifts *d* are enumerated within a predefined range [*−D, D*]. The shift *d* specifies the putative displacement of the second model relative to the first. If an anchor of the first model is located at position *p*, the corresponding position of the second model is considered to be *p* + *d*. Local windows of the same radius *R* are extracted around these two positions.

The same procedure is then performed in the opposite direction, starting from anchor positions of the second model. In this direction, the expected position of the first model can be refined within a small local neighborhood by selecting the maximum of its normalized profile. This accounts for small differences in the position of the local peak between the two models and reduces sensitivity to shifts of a few nucleotides. Window pairs obtained from the two directions are then pooled, and duplicates are removed.

#### Final profile comparison

For each orientation and shift, MIMOSA obtains aligned local-window collections for the two motif models. These collections are compared using the selected similarity metric, Dice (Costa, 2022) or cosine (Costa, 2022; Levy et al., 2025) (see below). The metrics are applied to continuous normalized values within windows. Therefore, the final score accounts not only for coordinate overlap of strong signals but also for similarity in the local profile structure around them.

The final similarity score for two motif models is defined as the maximum value over all considered orientations and shifts:

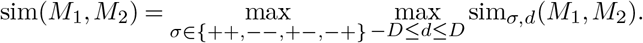

### E. Profile similarity metrics

In this work, we used two MIMOSA metrics for comparing normalized profiles: cosine similarity (Costa, 2022; Levy et al., 2025) and Dice similarity (Costa, 2022).

#### Cosine similarity

The cosine metric evaluates similarity in the shape of local profiles. It is computed separately for each pair of aligned windows, and the final value is obtained by averaging over all finite window-level scores:

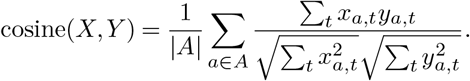

Here, *a* denotes an aligned window pair, *t* denotes a position within the window, and *A* is the set of windows for which both vector norms are non-zero. This metric characterizes not only the joint presence of strong signals but also their relative distribution within a local window. Therefore, cosine more strongly penalizes differences in profile shape, such as shifted maxima within the window or differences in the shape and width of local recognition profiles, even when both models mark a nearby sequence region.

#### Dice similarity

The dice metric measures the degree of symmetric overlap between positive signals in two profiles. Let *X* and *Y* denote two collections of aligned windows extracted from the profiles of two models after selecting the optimal shift and orientation. In the MIMOSA implementation, Dice similarity is first computed separately for each aligned window and then averaged across all windows, analogously to cosine similarity:

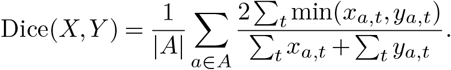

Here, *A* denotes the set of aligned windows, and the index *t* runs over positions within window *a*. If the total signal of both profiles in a window is close to zero, the local Dice value for that window is undefined and is excluded from averaging.

For non-negative values, the metric ranges from 0 to 1. A high value indicates that strong signals in the two profiles are well localized to the same positions within individual windows and have comparable mass.

### F. Statistical significance estimation

A critical component of motif comparison is statistical significance estimation, that is, determining whether the observed similarity could plausibly arise when unrelated motifs are compared. In MIMOSA, the null hypothesis is defined at the level of a motif collection with known group labels, such as transcription factor families or classes. In this work, we constructed the null distribution from HOCOMOCO motifs after removing motifs assigned to the unrecognized class and redundant motifs for the same TF. The remaining motifs were grouped according to TF class. For each motif, we generated a shuffled version by randomly permuting motif positions and then permuting nucleotide weights within positions. We then used the shuffled motifs to form null query–target comparisons: each eligible shuffled query motif was compared with shuffled target motifs from different TF classes; motifs from the same class were not used as null targets for each other.

Let *G*(*m*) denote the group label of motif *m*, and let *s*(*q, t*) be the value of the selected similarity metric for motifs *q* and *t* after optimization over shifts and orientations. The pooled null sample is then defined as

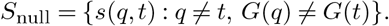

Thus, all eligible between-group comparisons contribute to a single shared null sample. This construction estimates the distribution of similarity expected for motifs not assigned to the same group and avoids fitting a separate null model for each query–target pair.

Because all reported metrics are oriented so that larger values indicate stronger motif similarity, the significance of an observed score *S*_obs_ is evaluated as an upper-tail probability:

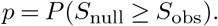

To obtain a smooth tail estimate, the pooled null distribution is approximated by a generalized extreme value distribution. Let *F*_GEV_(*x*; *θ*) and *f*_GEV_(*x*; *θ*) denote the cumulative distribution function and density of this distribution. Its parameters are estimated from *S*_null_, and the significance estimate is computed from the survival function:

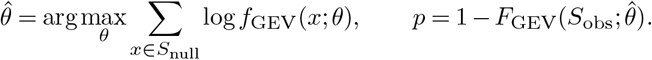

When multiple motif comparisons are annotated, the resulting P-values are further adjusted to control the false discovery rate, and the expected number of random matches is estimated as the product of the P-value and the effective number of target motifs.

### G. Implementation

MIMOSA is implemented in Python 3 and uses NumPy, Numba, and SciPy for numerical computation. The modular architecture separates model loading, profile generation, comparison, and statistical testing into distinct components, facilitating extension to new model types and similarity metrics. The software is designed for efficient large-scale analyses and supports batch processing of multiple motif comparisons.

## ACKNOWLEDGEMENTS

The research was funded by the Russian Science Foundation (project No. 25-74-00116).

## AUTHOR CONTRIBUTIONS

AT — data analysis, software development, and initial manuscript draft. VL — scientific supervision, data analysis.

## Supplementary Information

**Figure S1.**
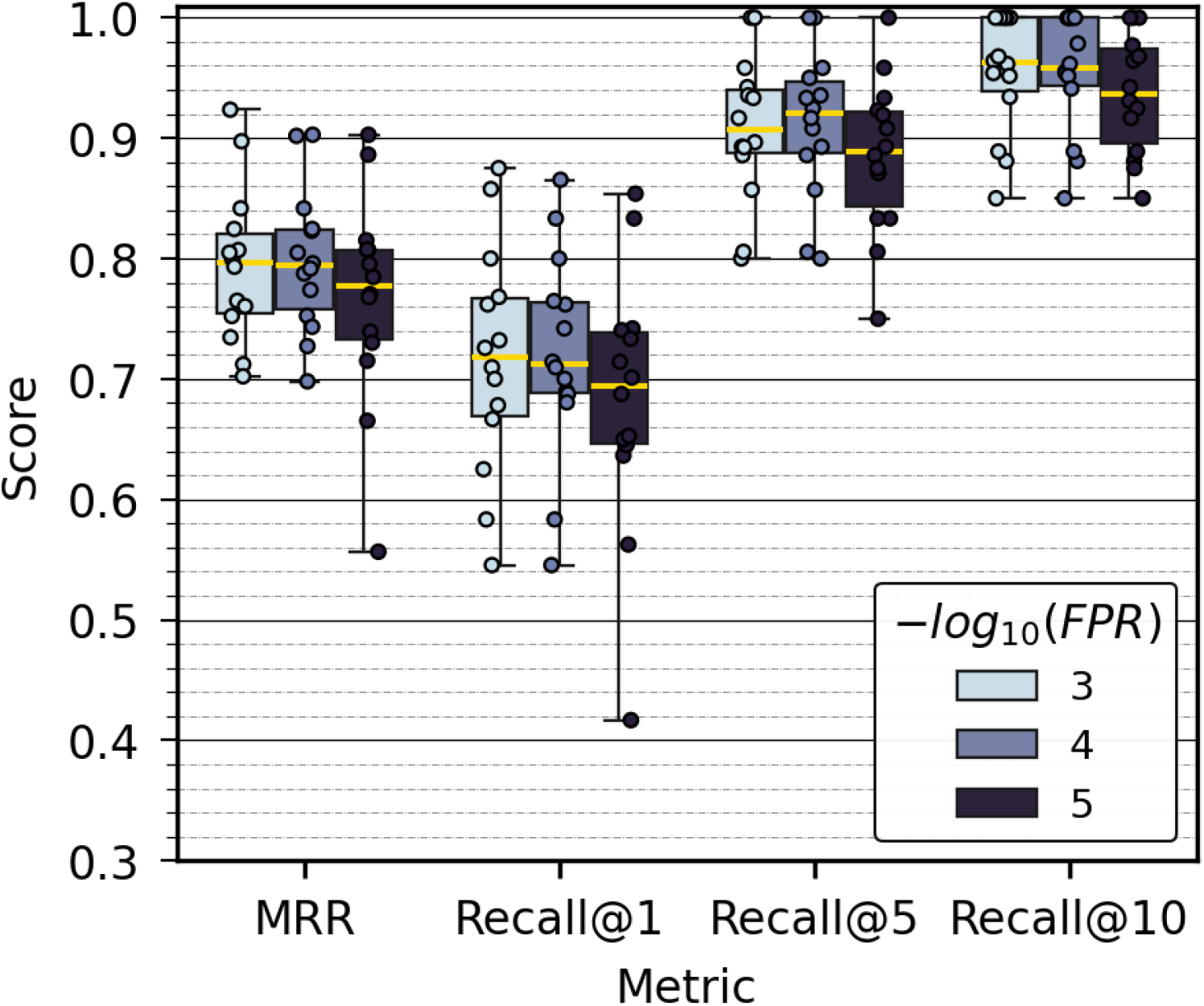
Accuracy assessment for mimosa-dice at different significance thresholds. Distribution of performance metrics for the MIMOSA variant using a Dice-like similarity measure for recognition profiles. The x-axis shows the ranking metrics: MRR, Recall@1, Recall@5, and Recall@10; the y-axis shows the corresponding metric value. Colors indicate the threshold on the allowed false-positive rate, expressed as −log_10_(FPR): 3, 4, and 5 correspond to FPR = 10^−3^, 10^−4^, and 10^−5^, respectively. Individual points show values for separate transcription factor classes, boxes indicate the interquartile range, horizontal yellow lines show the median, and whiskers indicate the range of observed values. Comparison across thresholds shows that ranking performance remains robust as the significance criterion becomes more stringent. The highest and most stable values are observed for Recall@5 and Recall@10, whereas Recall@1 is, as expected, the most stringent and variable metric.

**Figure S2.**
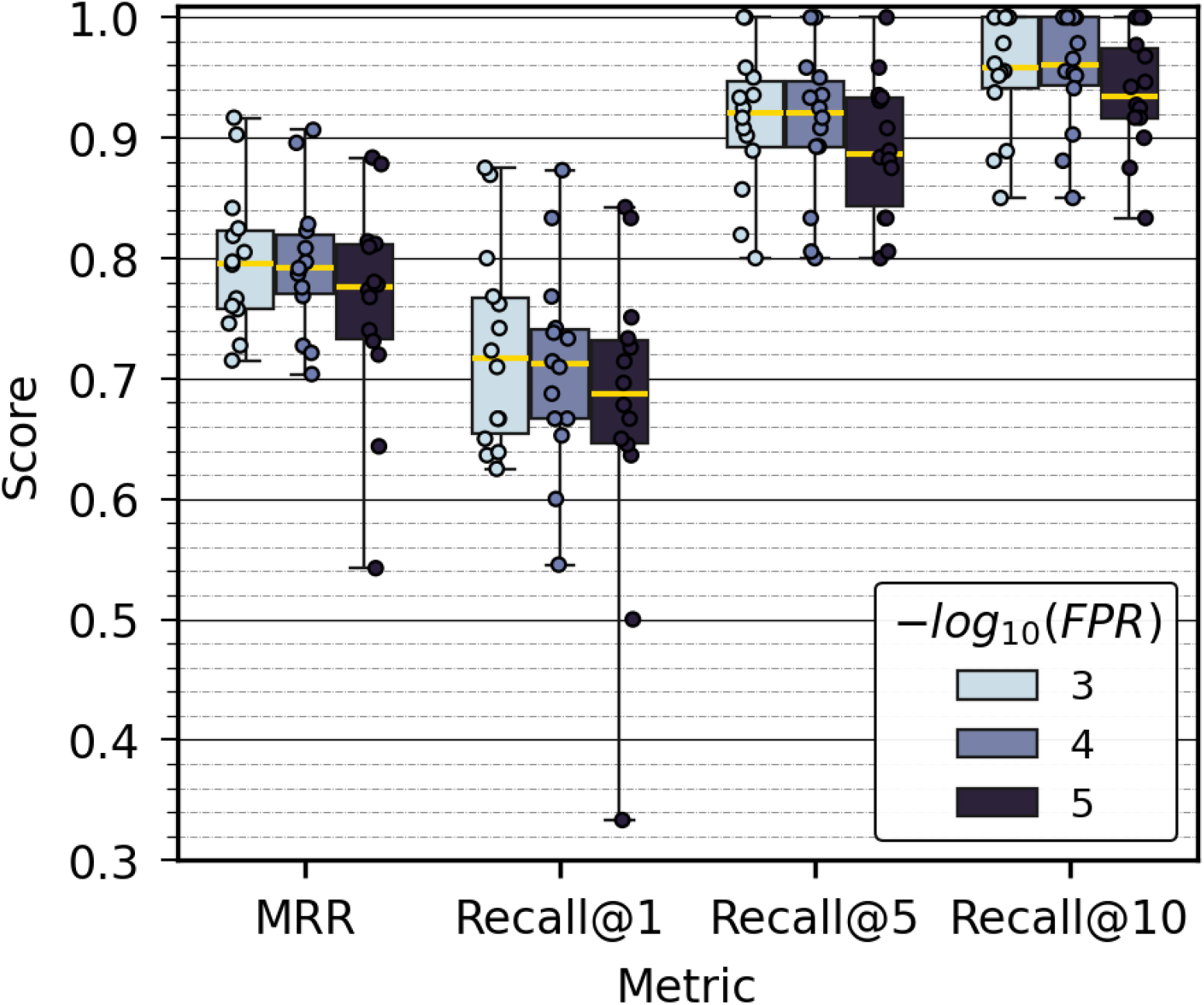
Accuracy assessment for mimosa-cosine at different significance thresholds. Distribution of MRR, Recall@1, Recall@5, and Recall@10 values for the MIMOSA variant based on cosine comparison of motif recognition profiles. Colors indicate three stringency levels for filtering matches by the false-positive rate: −log_10_(FPR) = 3, 4, and 5. Each point corresponds to the result for an individual transcription factor class, boxes show the interquartile range, yellow lines indicate medians, and whiskers show the range of observed values. As for the Dice-based variant, increasing the threshold stringency does not lead to a marked decrease in performance: Recall@5 and Recall@10 remain high for most classes, indicating that candidate rankings are robust to the choice of statistical threshold.

**Figure S3.**
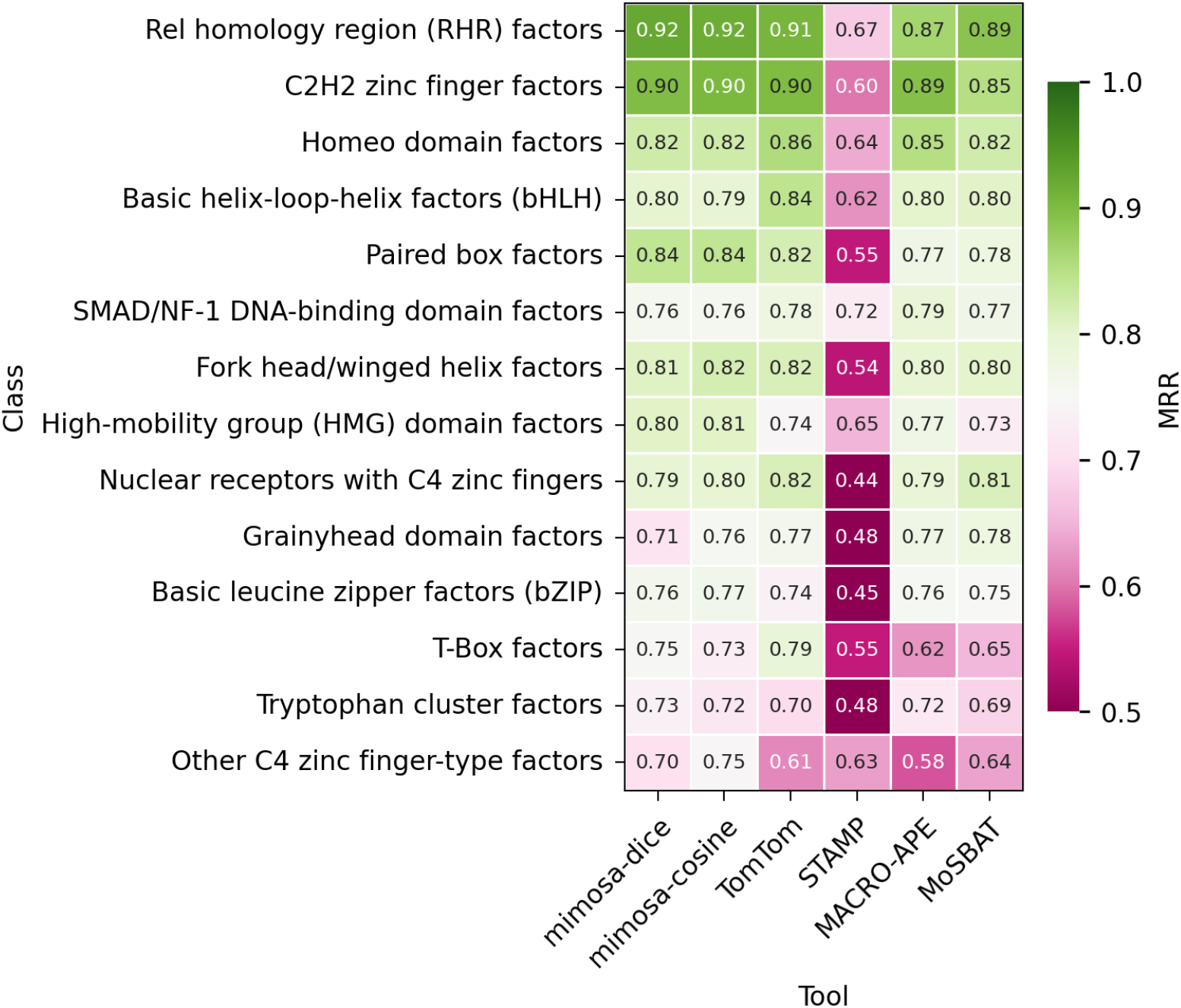
MRR by DNA-binding domain class. Heatmap of mean reciprocal rank (MRR) for different transcription factor classes and motif comparison tools. Rows correspond to structural classes of DNA-binding domains, and columns correspond to comparison methods: mimosa-dice, mimosa-cosine, Tomtom, STAMP, MACRO-APE, and MoSBAT. Each cell indicates how highly the first correct candidate is ranked for a given class; higher values correspond to better ranking and are highlighted in green. Both MIMOSA variants show robust MRR values across many classes, including RHR, C2H2 zinc finger, homeodomain, and bHLH factors. Lower values in individual rows indicate classes for which early ranking of the correct motif remains more challenging, such as T-box factors, tryptophan cluster factors, and some C4 zinc finger-type factors.

**Figure S4.**
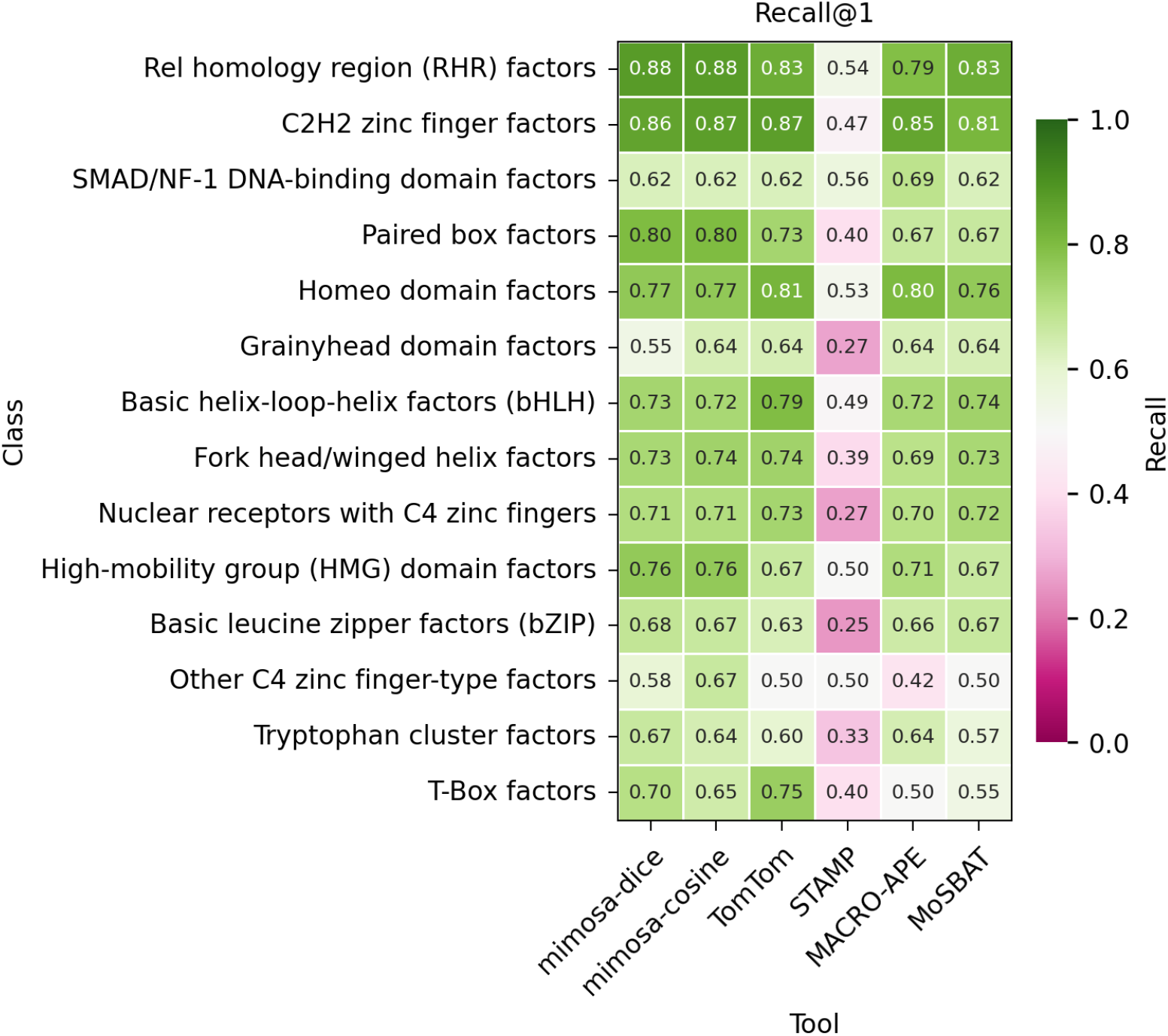
Recall@1 by DNA-binding domain class. Heatmap showing the fraction of queries for which the correct motif, or a motif from the correct class, was found at the first position of the ranked list. Rows denote transcription factor classes, and columns denote the comparison tools. The color scale represents Recall@1 values from 0 to 1: green cells indicate a higher probability that the top-ranked candidate is correct. This metric is most sensitive to errors in early ranking; therefore, differences between tools and classes are more pronounced here than for Recall@5 and Recall@10. The MIMOSA variants show high Recall@1 values for well-represented and structurally conserved classes, such as RHR and C2H2 zinc finger factors, whereas for some classes with more variable motifs, exact top-rank recovery is less stable.

**Figure S5.**
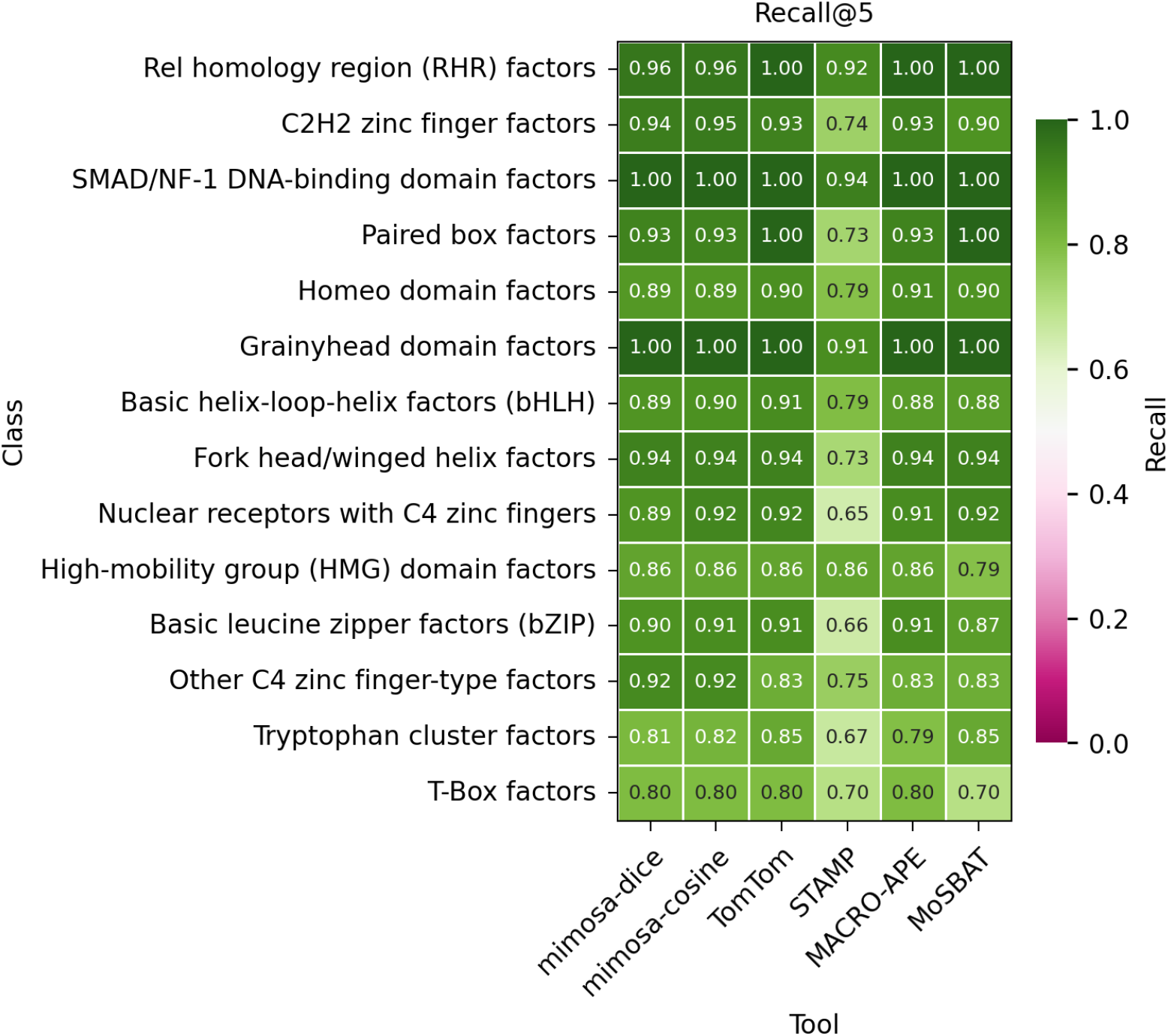
Recall@5 by DNA-binding domain class. Heatmap of Recall@5, showing the fraction of queries for which the correct motif is present among the five top-ranked results. Rows correspond to transcription factor classes, and columns correspond to motif comparison tools. Compared with Recall@1, this metric evaluates the practical utility of a method in a scenario where a researcher inspects several of the most similar database candidates. High Recall@5 values for most classes and methods show that the correct annotation usually appears near the top of the result list. The MIMOSA variants perform comparably to Tomtom, MACRO-APE, and MoSBAT and maintain high recall across a broad range of classes, including RHR, C2H2, SMAD/NF-1, paired box, forkhead/winged helix, and nuclear receptor motifs.

**Figure S6.**
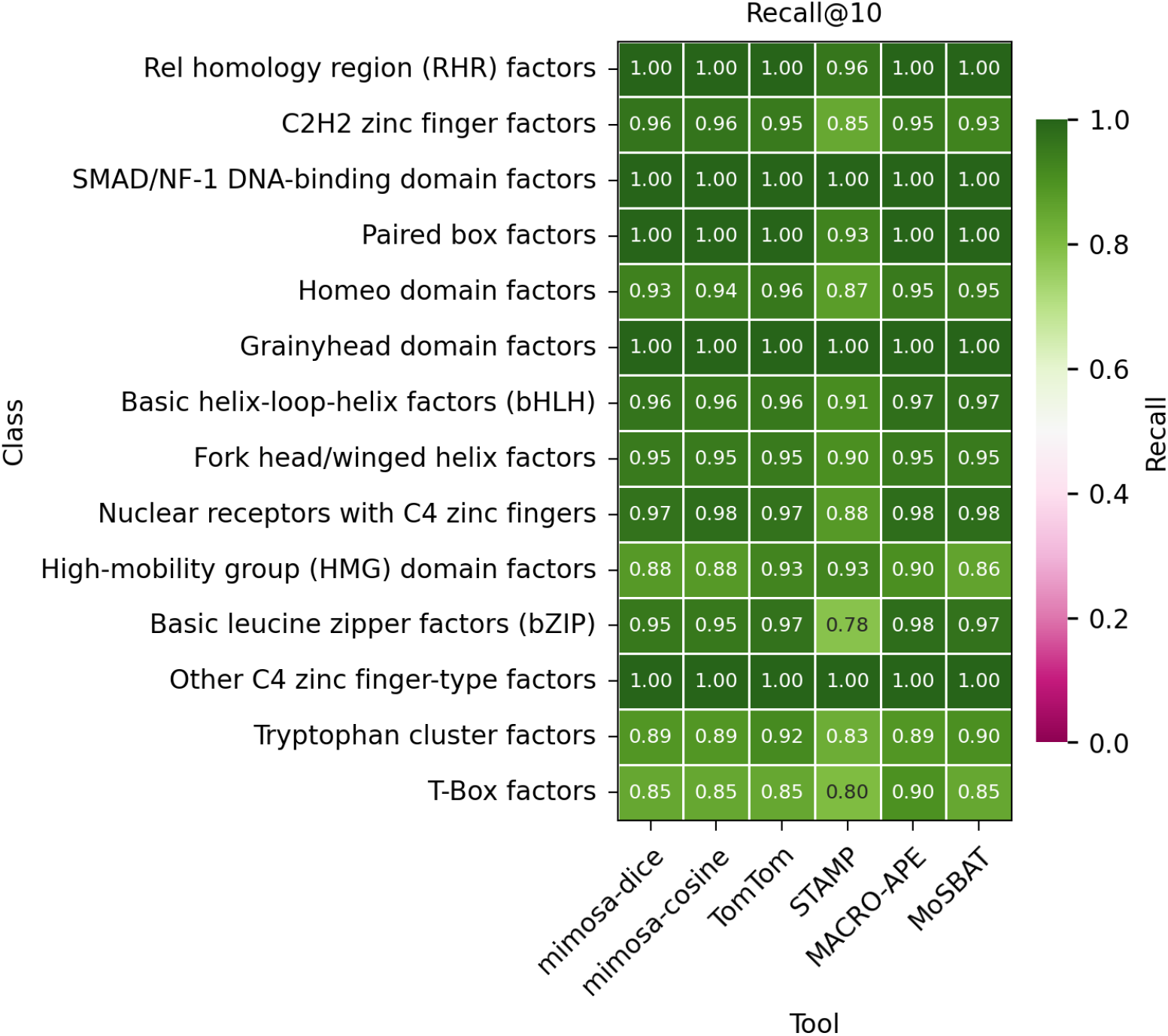
Recall@10 by DNA-binding domain class. Heatmap of Recall@10, showing the fraction of queries for which the correct motif is found among the ten top-ranked matches. This metric characterizes retrieval completeness when a broader candidate set is considered, which is relevant for downstream manual interpretation or automated motif annotation. Recall@10 values are high for almost all classes and reach 1.0 for several classes, indicating recovery of the correct match for all queries in those classes. These results show that even when the correct motif does not always occupy the first rank, most tools, including both MIMOSA variants, retain the ability to place it near the top of the ranked list.

**Figure S7.**
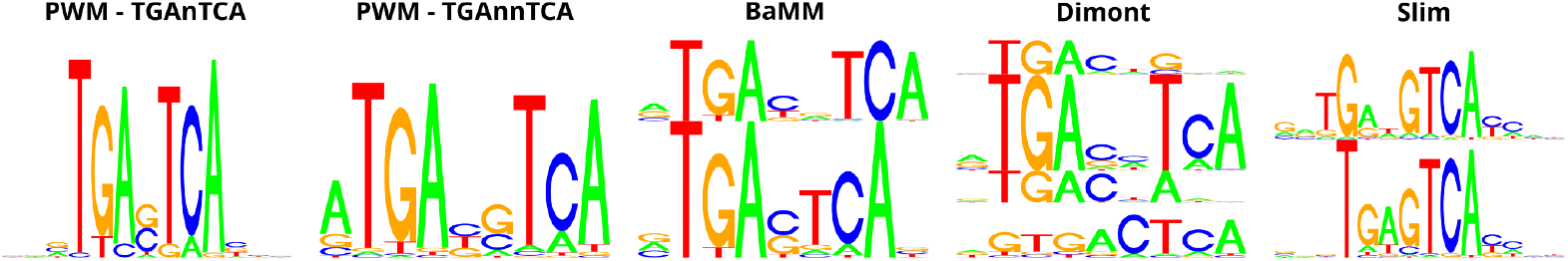
ATF3 site blocks generated with DepLogo. Sequence blocks are shown for two ATF3 PWM variants, TGAnTCA and TGAnnTCA, and for the BaMM, DIMONT, and Slim models. The block representation visualizes site heterogeneity within each motif. The PWM models separate variants that differ in spacer length into two distinct motifs, whereas BaMM and Slim contain blocks corresponding to both structural variants of the ATF3 site.

